# Descriptive epidemiology of energy expenditure in the UK: Findings from the National Diet and Nutrition Survey 2008 – 2015

**DOI:** 10.1101/542613

**Authors:** Soren Brage, Tim Lindsay, Michelle Venables, Katrien Wijndaele, Kate Westgate, David Collins, Les Bluck, Nick Wareham, Polly Page

## Abstract

**Background:** Little is known about population levels of energy expenditure as national surveillance systems typically employ only crude measures. The National Diet and Nutrition Survey (NDNS) in the UK measures energy expenditure in a 10% subsample by gold-standard doubly-labelled water (DLW).

**Methods:** DLW-subsample participants from the NDNS (383 males, 387 females) aged 4-91yrs were recruited between 2008 and 2015. Height and weight were measured, and bodyfat percentage was estimated by deuterium dilution.

**Results:** Absolute Total Energy Expenditure (TEE) increases steadily throughout childhood, ranging from 6.3 and 7.2 MJ/day in 4-7yr-old to 9.9 and 11.8 MJ/day for 14-16yr-old girls and boys, respectively. TEE peaked in 17-27yr-old women (10.9 MJ/day) and 28-43yr-old men (14.4 MJ/day), before decreasing gradually in old age. Physical Acitivty Energy Expenditure (PAEE) declines steadily with age from childhood (87.7 kJ/day/kg in 4-7yr olds) through to old age (38.9 kJ/day/kg in 71-91yr olds). Bodyfat percentage was strongly inversely associated with PAEE throughout life, irrespective of expressing PAEE relative to bodymass or fat-free mass. Compared to females with <30% bodyfat, females >40% recorded 28 kJ/day/kg and 17 kJ/day/kg fat-free mass less PAEE in analyses adjusted for age, geographical region, and time of assessment. Similarly, compared to males with <25% bodyfat, males >35% recorded 26 kJ/day/kg and 10 kJ/day/kg fat-free mass less PAEE.

**Conclusions:** This first nationally representative study reports levels of human energy expenditure as measured by gold-standard methodology; values may serve as reference for other population studies. Age, sex and body composition are main biological determinants of energy expenditure.

**Key messages:**

- First nationally representative study of human energy expenditure, covering the UK in the period 2008-2015
- Total Energy Expenditure (MJ/day) increases steadily with age thoughout childhood and adolescence, peaks in the 3^rd^ decade of life in women and 4^th^ decade of life in men, before decreasing gradually in old age
- Physical Acitivty Energy Expenditure (kJ/day/kg or kJ/day/kg fat-free mass) declines steadily with age from childhood to old age, more steeply so in males
- Bodyfat percentage is strongly inversely associated with physical activity energy expenditure

## Introduction

Little is known about population levels of energy expenditure (EE) as most national surveys use proxy methods for assessment, typically questionnaires. These may take the form of either self-reported dietary energy intake combined with measures of weight change^1^, or self-reported physical activity combined with estimates of resting EE^2^. The former approach is challenged not only by the necessary correction for any weight changes but also by possible underreporting of energy intake by overweight or obese individuals^3^. The latter approach does not need to make assumptions about energy balance as it is directly assessing the expenditure side; however, self-report methods for physical activity also have limited accuracy, and this applies particularly to derivatives such as estimates of energy expenditure^4^. The use of objective methods in the form of wearable sensors such as accelerometers and heart rate monitors is typically preferred as the objective methods for large-scale population studies, since these provide information about intensity patterns as well as more precise estimates of energy expenditure when coupled with appropriate inference models^5–8^. Irrespective of the success of such inference models, feasibility is somewhat limited for methods using heart rate monitoring due to its requirement for individual calibration using an exercise test ^9,10^, whereas the main limitation of accelerometry-based estimation of energy expenditure depends on the mix of specific behaviours in which the population under study is engaged as this relationship varies by activity type^11,12^.

Preferably, one would therefore employ more direct, yet highly feasible, measurements of the quantity of interest for the surveillance of population trends in energy expenditure. The doubly-labeled water (DLW) technique is the gold-standard for measurement of energy expenditure during free-living^13^. This technique uses the stable isotopes deuterium (^2^H) and Oxygen-18 (^18^O) to directly measure rate of carbon dioxide production (rCO_2_) over a period of 1-2 weeks, from which average total energy expenditure (TEE) can be calculated with high precision. Combined with simple anthropometric measurements, estimates of physical activity energy expenditure (PAEE) can also be derived. The DLW method is highly feasible in terms of low participant burden but it is unfortunately also expensive and hence is only seldom used in large studies.

The National Diet and Nutrition Survey (NDNS) employs a nationally representative sampling frame to assess dietary behaviours in the UK population^14^. One of the unique features of the NDNS is that a 10% subsample of all age groups 4years or older also had energy expenditure assessed using the DLW technique over 10 days of free-living. The aim of this study was to describe the variation in components of energy expenditure by key personal characteristics, geographical location, and over time.

## Methods

### Participants

Participants were recruited to the rolling programme in the main NDNS by stratified and clustered random sampling of households in the UK. NDNS data are weighted to account for any selection or response biases to ensure results are representative of the UK population^15^. A total of 15,583 households were selected to take part in the main NDNS, and 8,974 households agreed (58% household response rate). From those households, 10,727 individuals agreed to take part and a subsample of these main NDNS participants were invited to take part in the DLW substudy, within which individuals were sampled according to pre-specified age/sex strata (4-10, 11-15, 16-49, 50-64, and 65+ years). The DLW sub-study field work was carried out in two waves; for the first wave (2008-11), targets were 40 participants in each of the age/sex groups but for the second wave (2013-15), these were changed to 30 participants for each stratum for those aged 4-10 and those 65+ years, and to 50 participants for those aged 16-49 years. A total of 808 were invited to take part in the DLW substudy, of whom 770 participants provided sufficient data to derive valid EE estimates and they constitute the sample included in the present analysis. This subsample does not differ from the main NDNS (excluding children <4yrs) in terms of sex, body mass index (BMI), total energy intake, fruit and vegetable intake in g/day, free sugar intake (% total energy intake), and saturated fat intake (% total energy intake) but it was 2.6 years older^16^.

All adult participants provided informed written consent and all children provided assent with written consent from their legal guardian. The study was approved by the Oxfordshire A Research Ethics Committee (#07/H0604/113) and Cambridge South NRES Committee (#13/EE/0016).

### Measurements

Anthropometric measurements were performed in participants’ homes. Height was measured to the nearest millimeter using a portable stadiometer and bodymass was measured to the nearest 100g in light clothing using calibrated scales^17^. BMI (kg/m^2^) was calculated from these measures.

For the measurement of TEE, a baseline (pre-dose) urine sample was first collected to establish the natural abundance of the ^2^H and ^18^O isotopes in body water. Next, a dose of ^2^H_2_^18^O proportional to the participant’s bodymass (80mg per kg bodymass of ^2^H_2_O and 150mg per kg bodymass of H_2_^18^O) was prepared in a dose bottle. The full dose was drunk using a straw following which the bottle was re-filled with local tap water and again fully drunk by the participant. Participants collected single daily spot urine samples for the next ten days, representing about 2.5 half-lives of peak enrichment. The urine samples were analysed for isotopic enrichment by mass spectrometry (^18^O enrichment: AP2003, Analytical Precision Ltd, Northwich, Cheshire, UK; ^2^H enrichment: Isoprime, GV Instruments, Wythenshaw, Manchester, UK or Sercon ABCA-Hydra 20-22, Sercon Ltd, Crewe, UK). Rate of carbon dioxide production was measured using the method of Schoeller^18^ and converted to TEE using the energy equivalents of CO_2_ of Elia and Livesey^19^ assuming a respiratory exchange quotient of 0.85. Total bodywater was assessed using the zero-time intercept of deuterium turnover^20^ and fat-free bodymass calculated using a hydration factor of 73%^21^. Bodyfat percentage was calculated as total bodymass minus fat-free mass, expressed as percentage of total.

Resting metabolic rate was estimated from anthropometry variables by averaging three prediction equations; one based on age, sex, height, and total bodymass derived in a large database^22^, and two based on smaller studies which also take into account body composition^23,24^. In order to calculate 24-hour resting energy expenditure (REE), we integrated this resting metabolic rate value over time, but with a small adjustment for the 5% lower metabolic rate observed during sleep^25^ applied using age-specific sleep durations ranging from 8-12 hours/day^26^. The diet-induced thermogenesis (DIT) was assumed to constitute 10% of TEE^27^, and PAEE was calculated as the residual energy expenditure which sums with REE and DIT to make up TEE, according to the equation PAEE = TEE - REE - DIT = 0.9.TEE - REE.

### Statistics

We expressed daily TEE in absolute (MJ/day) units and both TEE and PAEE in relative units (kJ/day/kg bodymass). As sensitivity analyses, we also expressed energy expenditure in units scaled to fat-free bodymass and in alometrically-scaled units of kJ/day/kg^2/3^ bodymass, the latter based on the theoretical principle that absolute energy expenditure scales to bodily dimensions to the power of 2 and bodymass scales to bodily dimensions to the power of 3^28,29^. We present summary statistics (mean and standard deviation) of all estimates of energy expenditure by recruitment strata, ie age and sex groups. In addition, we present box plots (box denoting median and interquartile ranges) by expanded age-groups (deciles), as well as by survey year (2008-11 and 2012-15) and main geographical regions of North England, South England, and Scotland/Wales/North-Ireland combined. North England included the following Government Office Regions; North East, North West, Yorkshire and The Humber, East Midlands and West Midlands, and South England comprised the East, South West, London and South East as used previously^30^. We examine the association with obesity status by both body mass index (BMI) and bodyfat groups, stratified by sex and age groups. To examine independent associations, we performed a multiple linear regression analysis with mutual adjustment for all above factors, and with additional adjustment for season of measurement (expressed as two orthogonal sine functions; “winter” (with max=1 on January 1^st^ and min=−1 on July 1^st^) and “spring” (with max=1 on April 1^st^ and min=−1 on October 1^st^).

## Results

Of the 770 participants with valid DLW data in the NDNS included in this analysis, the four constituent countries of the United Kingdom were represented with 568 participants from England, 50 from Scotland, 72 from Wales and 80 from Northern Ireland (Table 1).

**Table 1.**
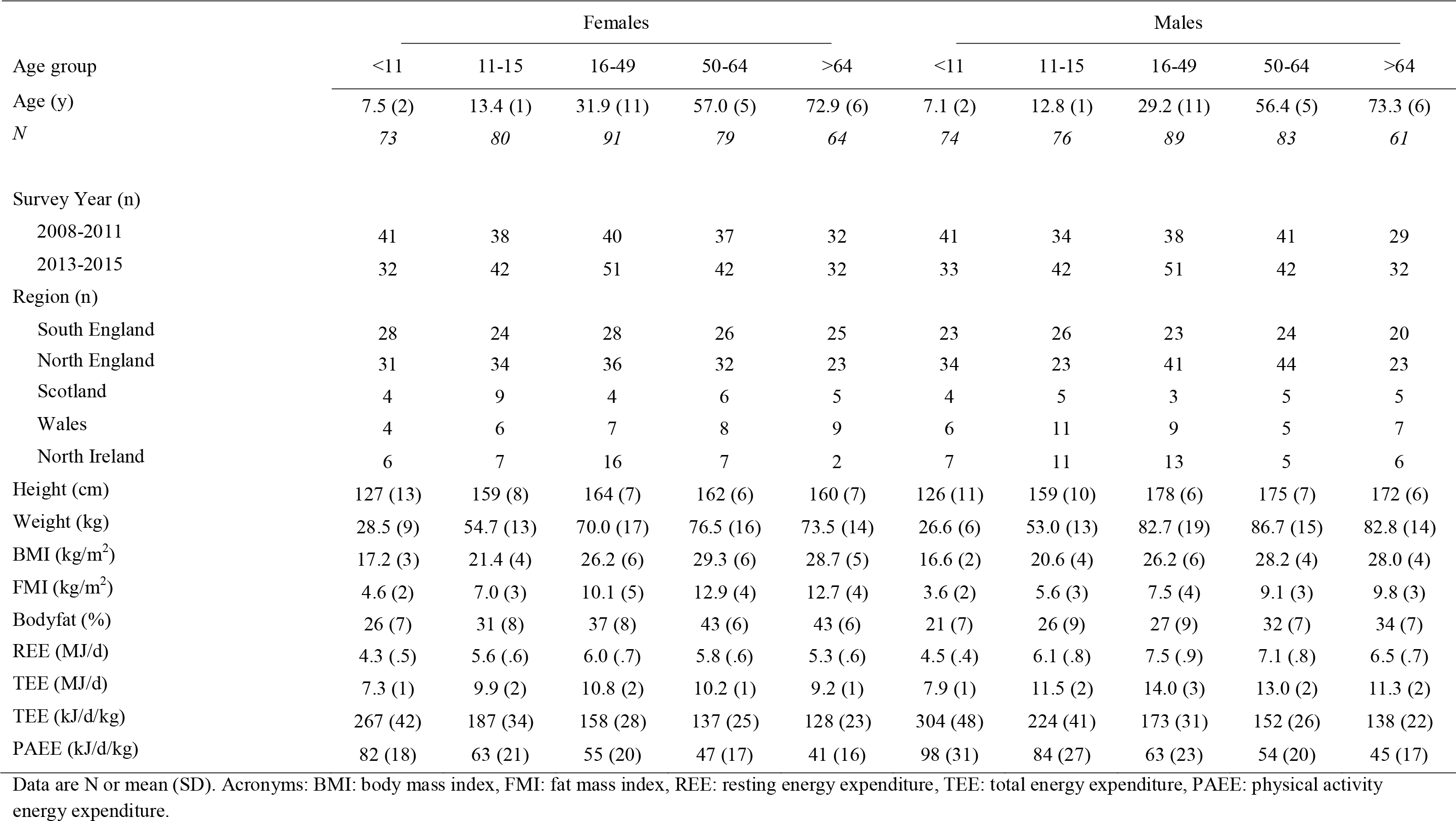
Participant characteristics. National Diet and Nutrition Survey DLW subsample (2008-2015)

Mean (SD) TEE was 10.6 (2.8) MJ•day^−1^ or 187 (64) kJ•day^−1^•kg^−1^, REE was 5.9 (1.2) MJ•day^−1^, and PAEE was 63 (28) kJ•day^−1^•kg^−1^ Across these estimates of energy expenditure, after adjustment for age, time and region of measurement, male sex was associated with higher values (p<0.001). When TEE and PAEE were expressed relative to fat-free mass, only PAEE was higher in males (p=0.007).

Figure 1 shows TEE, PAEE, and bodymass across age deciles and stratified by sex. Median TEE and PAEE were higher in males than females across all age groups; bodymass was similar in boys and girls up to age 16 yrs but higher in men above that age. Absolute TEE (MJ•day^−1^) was highest in 17-27-year old women and 28-43-year old men, respectively. In contrast, TEE and PAEE relative to bodymass (kJ•day^−1^•kg^−1^) was highest in the youngest individuals and displayed a consistent downward trend with advancing age into adulthood. TEE had a less steep association with age from early to later adulthood. The bodymass-scaled EE associations partially mirrored the positive trend in bodymass from childhood into young adulthood, which levelled off across adult ages. Similar age associations were observed in the sensitivity analyses (Supplementary Figure S1-S2), although the age association for allometrically scaled TEE (kJ•day^−1^•kg^−2/3^) was more linear across the whole age range, and 8-11-year olds had the highest PAEE of all groups in these analyses.

**Figure 1.**
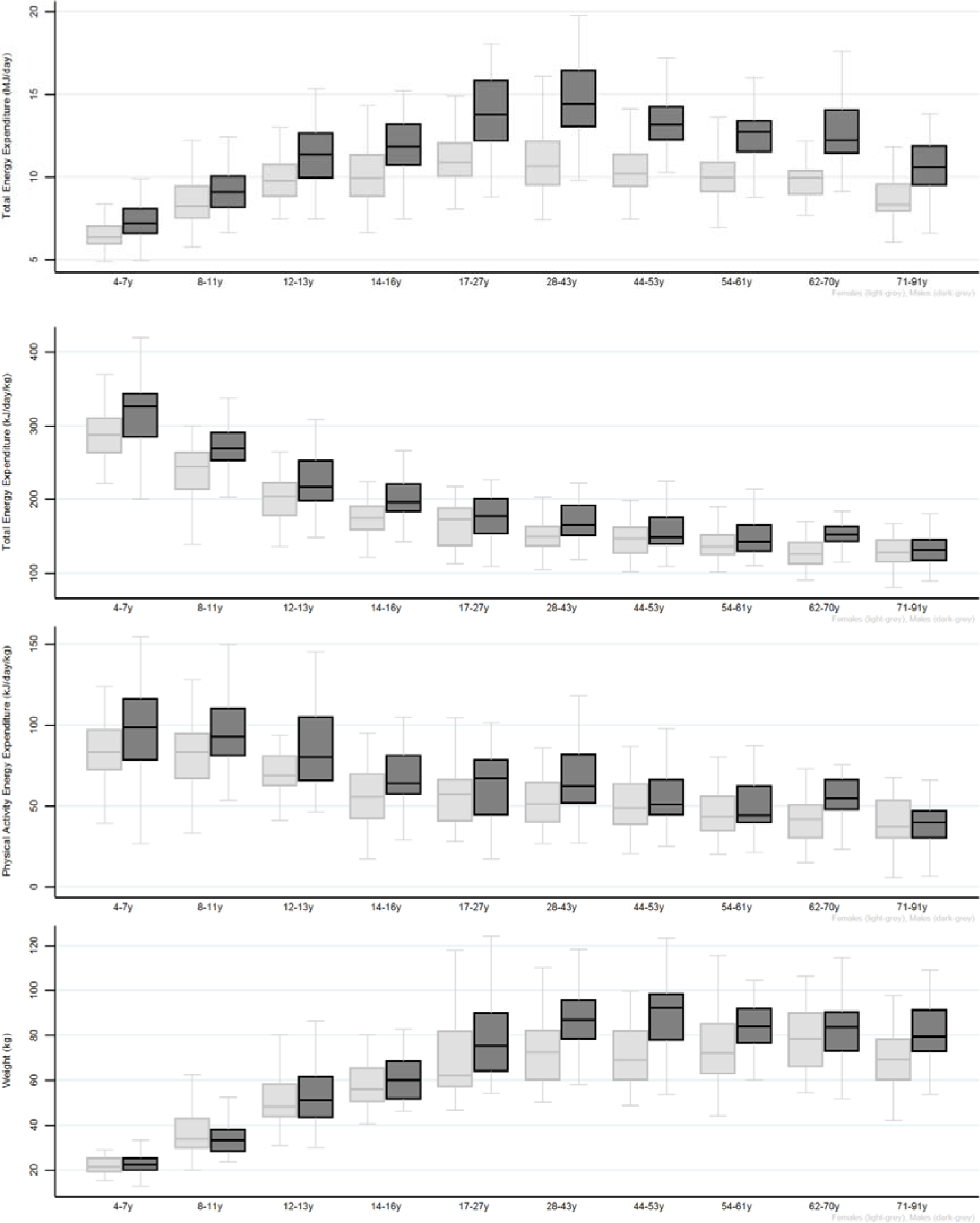
Total and Physical Activity-related Energy Expenditure by sex (Females= light grey, Males= dark grey) and age groups (approximate deciles). Bottom panel shows body weight by sex and age.

There were no significant differences in TEE, PAEE or bodymass among those participants surveyed between 2008-2011 and those surveyed between 2013-2015 (Figure 2), nor were there any discernible differences between constituent geographical regions (Figure 3). These observations were confirmed in the multi-variable adjusted analyses which were additionally adjusted for season of measurement, an effect which was only apparent in males, with slightly higher values in spring (Table 2).

**Table 2.** Mutually adjusted associations with energy expenditure. National Diet and Nutrition Survey DLW subsample (2008-2015)

| Women |  |  |  |  |  |  |
| --- | --- | --- | --- | --- | --- | --- |
|  | Total Energy Expenditure (MJ / day) | 95% C.I. | Total Energy Expenditure (kJ / day / kg) | 95% C.I. | Physical Activity Energy Expenditure (kJ / day / kg) | 95% C.I. |
| **Age** |  |  |  |  |  |  |
| 4-10y | Reference |  | Reference |  | Reference |  |
| 11-15y | 2.41*** | 1.92; 2.90 | -75.10*** | -83.84; -66.36 | -17.42*** | -23.13; -11.71 |
| 16-49y | 2.99*** | 2.50; 3.49 | -94.73*** | -103.56; -85.90 | -21.15*** | -26.92; -15.38 |
| 50-64y | 1.91*** | 1.35; 2.47 | -103.14*** | -113.13; -93.16 | -24.43*** | -30.95; -17.90 |
| 65-91y | 0.91*** | 0.34; 1.49 | -115.08*** | -125.32; -104.84 | -31.35*** | -38.04; -24.66 |
| **Year of Study** |  |  |  |  |  |  |
| 2008-2011 | Reference |  | Reference |  | Reference |  |
| 2013-2015 | 0.02 | -0.29; 0.33 | 3.47 | -2.05; 9.00 | 0.88 | -2.73; 4.49 |
| **Season** |  |  |  |  |  |  |
| Spring | 0.02 | -0.20; 0.24 | -0.03 | -3.90; 3.85 | 0.17 | -2.37; 2.70 |
| Winter | 0.11 | -0.11; 0.33 | 0.38 | -3.60; 4.36 | 1.19 | -1.41; 3.79 |
| **Region** |  |  |  |  |  |  |
| South England | Reference |  | Reference |  | Reference |  |
| North England | -0.07 | -0.43; 0.29 | -3.03 | -9.36; 3.29 | -1.45 | -5.58; 2.68 |
| Scotland, Wales, Northern Ireland | 0.18 | -0.22; 0.58 | 5.36 | -1.77; 12.49 | 3.84 | -0.82; 8.50 |
| **BMI Category** |  |  |  |  |  |  |
| <25 kg/m<sup>2</sup> | Reference |  | Reference |  | Reference |  |
| 25-30 kg/m<sup>2</sup> | 0.75*** | 0.32; 1.18 | -30.12*** | -37.75; -22.49 | -12.30*** | -17.29; -7.31 |
| >30 kg/m<sup>2</sup> | 1.79*** | 1.34; 2.25 | -43.25*** | -51.34; -35.15 | -17.86*** | -23.15; -12.58 |
| Constant | 7.28*** | 6.86; 7.70 | 266.65*** | 259.14; 274.16 | 81.89*** | 76.98; 86.79 |
| **<u>Model 2 (BF% instead of FMI category)</u>** |  |  |  |  |  |  |
| F: <30% M: <25% | Reference |  | Reference |  | Reference |  |
| F: 30-40% M: 25-35% | 0.09 | -0.39; 0.57 | -31.66*** | -38.81; -24.51 | -14.12*** | -19.02; -9.22 |
| F: >40% M: >35% | 0.72*** | 0.21; 1.23 | -61.72*** | -69.35; -54.08 | -28.20*** | -33.43; -22.98 |
| **Men** |  |  |  |  |  |  |

|  | Total Energy<br>Expenditure<br>(MJ / day) | 95% C.I. | Total Energy<br>Expenditure<br>(kJ / day / kg) | 95% C.I. | Physical<br>Activity Energy<br>Expenditure<br>(kJ / day / kg) | 95% C.I. |
| --- | --- | --- | --- | --- | --- | --- |
| **Age** |  |  |  |  |  |  |
| 4-10y | Reference |  | Reference |  | Reference |  |
| 11-15y | 3.34*** | 2.70; 3.98 | -75.57*** | -86.28; -64.87 | -12.94*** | -20.75; -5.14 |
| 16-49y | 4.97*** | 4.29; 5.64 | -111.24*** | -122.51; -99.97 | -27.17*** | -35.39; -18.96 |
| 50-64y | 3.42*** | 2.68; 4.17 | -126.35*** | -138.82; -113.88 | -34.32*** | -43.41; -25.23 |
| 65-91y | 1.64*** | 0.83; 2.44 | -139.98*** | -153.46; -126.50 | -43.27*** | -53.10; -33.45 |
| **Year of Study** |  |  |  |  |  |  |
| 2008-2011 | Reference |  | Reference |  | Reference |  |
| 2013-2015 | -0.16 | -0.57; 0.25 | -4.19 | -11.06; 2.67 | -2.88 | -7.88; 2.12 |
| **Season** |  |  |  |  |  |  |
| Spring | 0.38*** | 0.11; 0.66 | 2.46 | -2.15; 7.07 | 3.03* | -0.32; 6.39 |
| Winter | 0.20 | -0.10; 0.50 | 0.03 | -4.96; 5.02 | -0.08 | -3.72; 3.56 |
| **Region** |  |  |  |  |  |  |
| South England | Reference |  | Reference |  | Reference |  |
| North England | 0.18 | -0.29; 0.65 | -1.59 | -9.45; 6.26 | 0.20 | -5.53; 5.92 |
| Scotland, Wales, Northern Ireland | 0.21 | -0.33; 0.74 | 3.33 | -5.60; 12.26 | 4.25 | -2.26; 10.76 |
| **BMI Category** |  |  |  |  |  |  |
| <25 kg/m<sup>2</sup> | Reference |  | Reference |  | Reference |  |
| 25-30 kg/m<sup>2</sup> | 1.47*** | 0.91; 2.03 | -28.36*** | -37.75; -18.96 | -10.81*** | -17.66; -3.96 |
| >30 kg/m<sup>2</sup> | 2.88*** | 2.24; 3.53 | -41.05*** | -51.84; -30.26 | -15.50*** | -23.37; -7.64 |
| Constant | 7.87*** | 7.31; 8.43 | 306.51*** | 297.15; 315.87 | 98.54*** | 91.72; 105.37 |
| **Model 2 (BF% instead of FMI category)** |  |  |  |  |  |  |
| F: <30% M: <25% | Reference |  | Reference |  | Reference |  |
| F: 30-40% M: 25-35% | 0.54** | 0.00; 1.08 | -37.07*** | -44.38; -29.75 | -16.33*** | -22.01; -10.64 |
| F: >40% M: >35% | 0.86*** | 0.23; 1.48 | -55.68*** | -64.20; -47.16 | -25.88*** | -32.50; -19.25 |

C.I: confidence intervals
\*\*\* p<0.01, \*\* p<0.05, \* p<0.1

**Figure 2.**
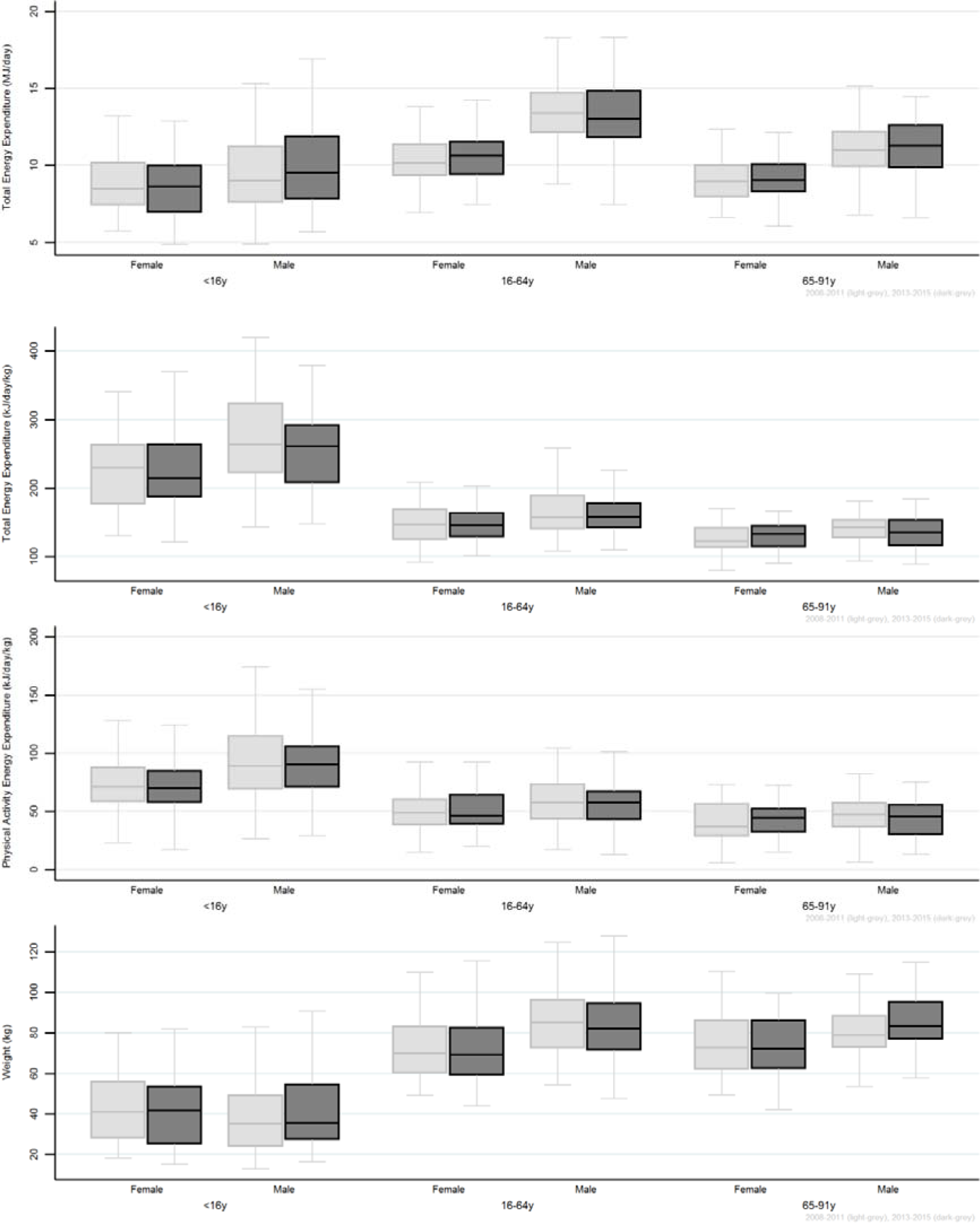
Age and sex-specific Total and Physical Activity-related Energy Expenditure by survey year (2008-2011= light grey, 2013-2015= dark grey). Bottom panel shows stratified body weight.

**Figure 3.**
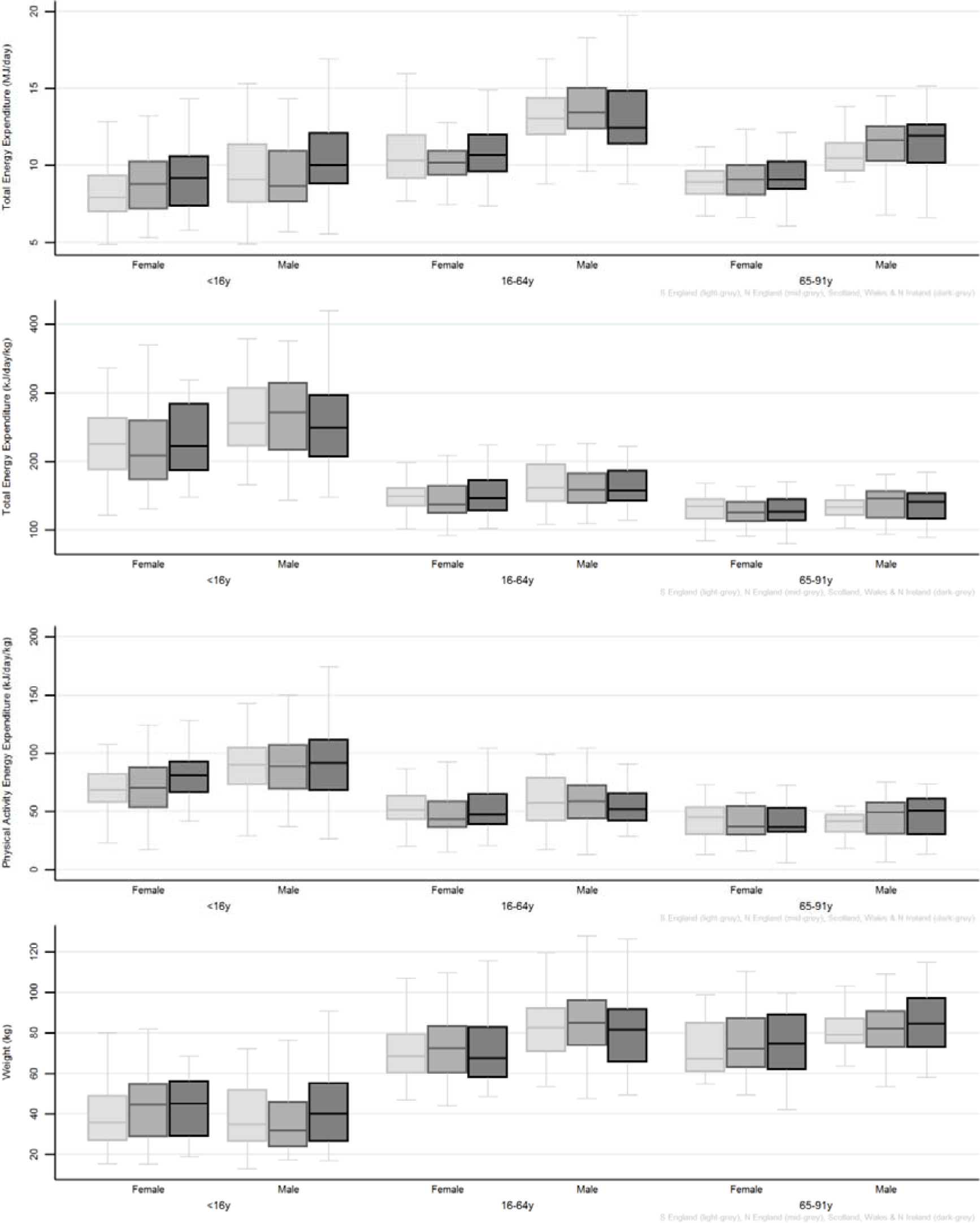
Age and sex-specific Total and Physical Activity-related Energy Expenditure by geographical region (South England = light grey, North England = medium-grey, Scotland, Wales, and North Ireland = dark-grey). Bottom panel shows stratified body weight.

Across the sample, absolute TEE (MJ•day^−1^) was higher in individuals with higher BMI. Overweight participants had higher TEE (MJ•day^−1^) than normal-weight participants, and obese participants accumulated higher TEE levels than overweight participants, a trend that was observed within nearly all age- and sex strata (Figure 4). However, this relationship was inverse when TEE was expressed in relative terms. Obese males and females in all age groups recorded the lowest relative TEE and PAEE (kJ•day^−1^•kg^−1^), whereas normal-weight individuals recorded the highest.

**Figure 4.**
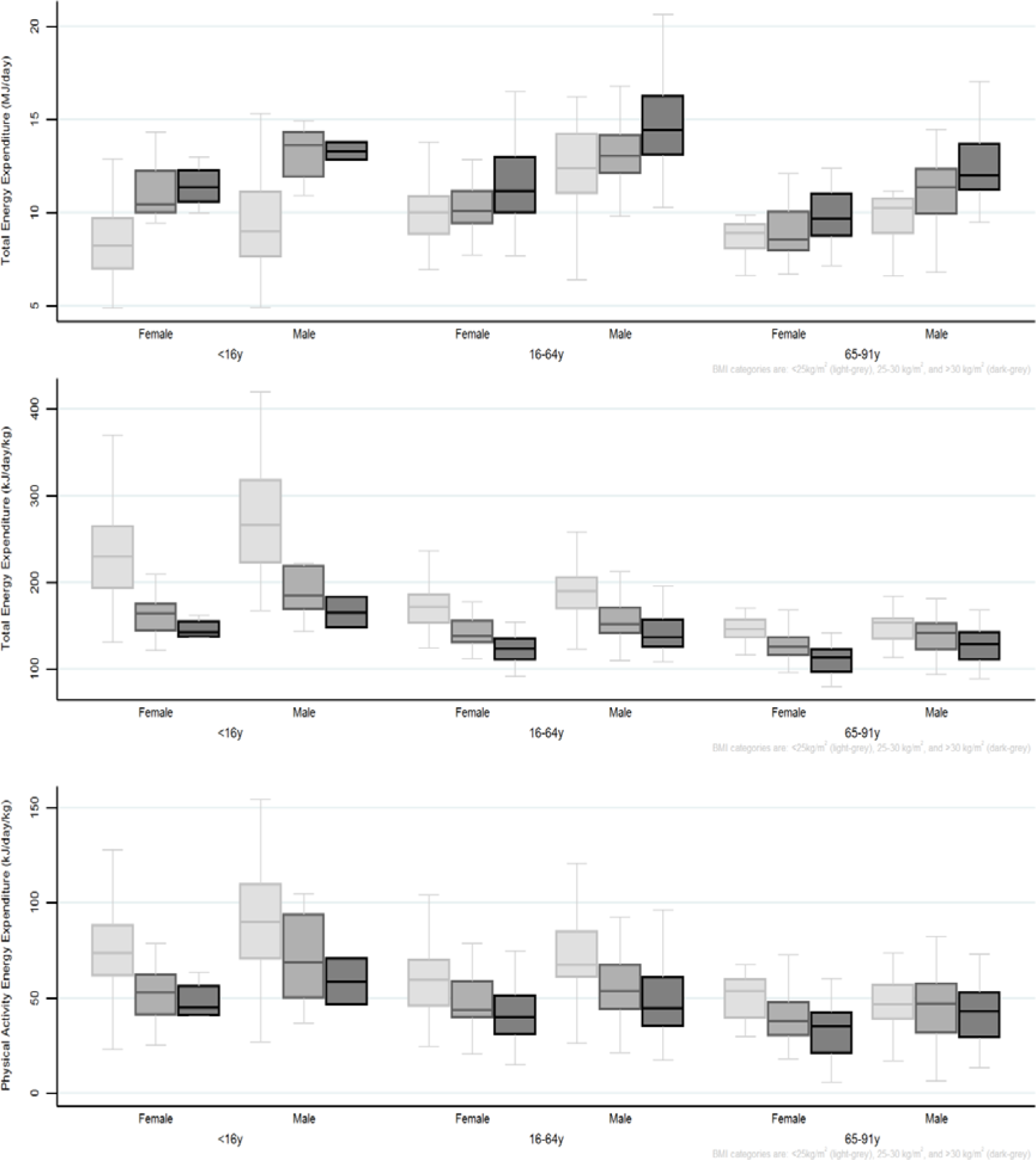
Age and sex-specific Total and Physical Activity-related Energy Expenditure by BMI category (Normal-weight (<25kg/m^2^) = light grey, Overweight (25-30kg/m^2^) = medium grey, Obese (>30kg/m^2^)= dark grey).

A similar relationship was also observed for TEE and PAEE across groups of differing bodyfat percentage, although the clear positive trend for absolute TEE was absent in the two adult age groups (Figure 5). For relative TEE and PAEE (kJ•day^−1^•kg^−1^), those with the highest bodyfat percentage recorded the lowest energy expenditure, whereas the slimmest individuals recorded the highest. The sole exception to this were men aged 65-91y with medium bodyfat who as a group accumulated more PAEE than their slimmer counterparts. The multivariable regression analysis confirmed associations with BMI and body composition in both sexes (Table 2).

**Figure 5.**
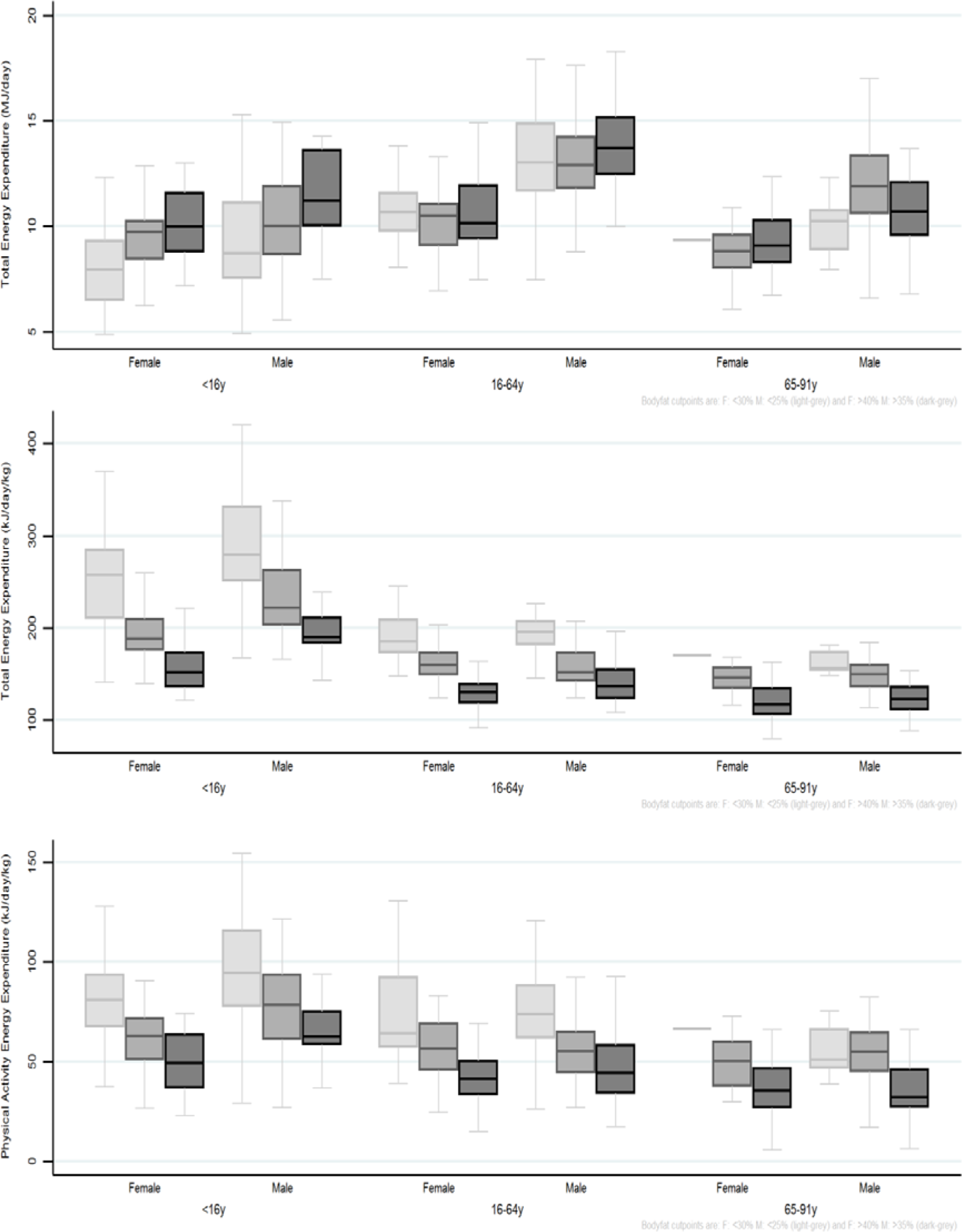
Age and sex-specific Total and Physical Activity-related Energy Expenditure by bodyfat% (Slimmest = light grey, medium body composition = medium grey, fattest = dark grey).

In sensitivity analyses modelling PAEE per kg fat-free mass (supplement table T2), individuals in the third tertile of fat mass index were less active; this inverse association was more consistent for bodyfat percentage groups. This sensitivity analysis also suggested a possible regional difference in activity levels, with non-English women expending more activity energy per kg fat-free mass, independent of other covariates.

## Discussion

Here, we report gold-standard measured energy expenditure from a nationally representative UK survey. Our results show that TEE and PAEE vary according to age, sex and body composition but not significantly so by geographical region of the UK or over time in the period between 2008 and 2015.

Our results demonstrate that males accumulate higher overall levels of TEE and PAEE than females across all ages, a finding that is consistent with other British cohort studies investigating energy expenditure with objective methods^31–34^. Age was an important correlate of PAEE and TEE in both sexes, with similar patterns across the lifespan for all EE measures; absolute TEE peaks in the early adult years, before dropping off around retirement age, whereas relative TEE and PAEE are highest in the earliest years of life before gradually declining steeply at first and reflecting in part natural growth and development, and then more shallowly after the age when adult height is typically attained.

To our knowledge no large DLW studies exist in paediatric populations; participants in the NDNS recorded similar levels of PAEE (93 vs 95 kJ•day^−1^•kg^−1^) to 1397 age-matched, British 6-yr olds as measured by combined heart rate and movement sensing^31^, in fact several British cohort studies using this technique observe comparable PAEE levels across the age range, with 74 kJ•day^−1^•kg^−1^ in 825 adolescents (aged 15 years) attending schools in Cambridge^32^ and 54 kJ•day^−1^•kg^−1^ in a sample of 5442 English adults aged between 29 and 62 years (mean 48 yrs)^35^ and 34 kJ•day^−1^•kg^−1^ in 1787 members of the 1946 birth cohort assessed at ages 60-64 yrs^34^. Smaller DLW studies in children in developed countries have been reported; one study in 12 Australian toddlers from 2010 reported lower levels of TEE (5.2 MJ•day^−1^) than 4-7-year olds in the NDNS (6.9 MJ•day^−1^) but were also younger (3.2 years)^24^. Comparatively low TEE and PAEE were recorded at 4.8 MJ•day^−1^ and 44 kJ•day^−1^•kg^−1^ in a study of 97 healthy Texan children aged 4.5years^36^.

A Danish study in 26 boys and girls aged 9 years in 1999 reported TEE of 8.9 and 8.3 MJ•day^−1^, respectively^37^, compared to 9.3 and 8.5 MJ•day^−1^ in 8-11 year olds in NDNS; however PAEE was lower in the Danish study, 86 and 64 kJ•day^−1^•kg^−1^. A Swedish DLW study reported TEE of 13.8 and 10.7 MJ•day^−1^ in fifty 15-year old boys and girls, compared to 12.1 and 10.0 MJ•day^−1^ for the 14-16-year age group in NDNS^38^; PAEE was also higher (107 and 81 kJ•day^−1^•kg^−1^). Combined, these studies show similar absolute TEE in similarly aged children but varying levels of relative PAEE, although direct comparisons should bear in mind notable differences between studies, including participant selection, setting, and era.

In adults, male NDNS participants accumulated a mean TEE and PAEE of 12.9 MJ•day^−1^ and 54 kJ•day^−1^•kg^−1^, whereas adult women accumulated 10.1 MJ•day^−1^ and 47 kJ•day^−1^•kg^−1^, respectively. This is comparable to levels of TEE and PAEE in other DLW studies in comparable populations, e.g. mean TEE of 12.7 MJ•day^−1^ for men and 10.0 MJ•day^−1^ for women, and PAEE of 54 kJ•day^−1^•kg^−1^ and 44 kJ•day^−1^•kg^−1^ were reported in a meta-analysis of 1575 men and 2914 women aged over 19 years from high-development index countries^39^. This analysis included published DLW data upto 2011, and although studies in special populations were excluded, again caution is warranted as to the representativeness of the participants included.

More recently, Matthews et al reported DLW results from a study of 461 American men and 471 women in a convenience sample with mean (SD) ages of 64 (6) and 62 (6) years, respectively. In that study, mean TEE was 11.6 and 9.1 MJ•day^−1^ and mean PAEE was 39.2 and 37.5 kJ•day^−1^•kg^−1^, for men and women respectively^40^. Again, these figures are very similar to those found for TEE in the oldest category of NDNS participants (>64y), and only about 10% lower for PAEE although we note mean age was 72 years in our sample. Overall, the results therefore suggest that British men and women expend a similar amount of total and physical activity energy as their counterparts in the developed world, with a similar age-related decline.

This is partially in contrast to EE levels in populations residing in less developed countries, where only absolute EE levels are similar but activity levels are higher. For example, absolute TEE in studies from countries with low-to-medium development scores was reported to be 12.3 and 9.3 MJ/day but relative PAEE estimated at 68.5 and 48.9 kJ•day^−1^•kg^−1^, for men and women respectively^39^. With respect to PAEE, these high levels seem particularly pronounced in rural dwellers in these countries, with values around 60 kJ•day^−1^•kg^−1^ in Cameroon^41^ and even higher in rural Luo, Kamba, and Masai in Kenya^42^ as assessed with individually calibrated combined heart rate and movement sensing.

BMI and bodyfat percentage were also important correlates of TEE and PAEE, and there is an ongoing debate over how to best express energy expenditure with respect to body size, particularly when examining associations with overweight and obesity^43,44^ In the NDNS sample, larger body size was associated with higher absolute levels of TEE (MJ•day^−1^) but irrespective of how energy expenditure was expressed relative to body weight (kJ•day^−1^•kg^−1^ or kJ•day^−1^•kg^−2/3^), BMI displayed an inverse relationship. This was also observed when body composition was assessed in terms of total bodyfat % or fat mass index. Multivariate analysis demonstrated that, when corrected for age, geographical region, survey year and season of measurement, overweight women accumulated 31 kJ•day^−1^•kg^−1^ less TEE and 13 kJ•day^−1^•kg^−1^ less PAEE than their normal-BMI counterparts. Continuing this trend, obese females accumulated 44 kJ•day^−1^•kg^−1^ less TEE and 18 kJ•day^−1^•kg^−1^ less PAEE than normal-weight female participants. This was replicated in males with TEE and PAEE lower in groups with higher BMI.

This finding highlights the role that absolute body size plays in the accumulation of absolute energy expenditure, but also underlines obesity’s inverse association with physical activity energy expenditure. This relationship was apparent regardless of the measure of obesity and of the measure of physical activity, with those with higher absolute bodyfat levels and those in the highest FMI category accumulating less physical activity than slimmer counterparts.

This study has several notable strengths. Firstly, the NDNS is nationally representative and with no observed selection bias for the DLW subsample; therefore the estimates for TEE and PAEE can serve as national reference values for this period. Secondly, DLW is the gold standard method for measuring energy expenditure during free-living conditions. Thirdly, our analyses include the main components and common expressions of energy expenditure, including both absolute and various relative measures, and within the limitations of the sample also reasonable stratification and multivariable adjustment analyses to test the robustness of observed differences across specific population subgroups.

This study also has some limitations. The study as a whole is not large, with only 770 individuals included in the present analyses. In addition, the majority of the sample came from England, with very few participants included in certain subgroup analyses. The generalisability of these small groups to the wider Northern Irish, Scottish and Welsh populations is therefore less certain. It is also possible that non-participating households may differ from participating households. Another limitation is that data are effectively snap-shot assessments taken at relatively short time intervals between the 2008 and 2015 which is unlikely to be sufficient to detect secular trends even if they truly occurred in the UK over this time period; given the slight increase in national obesity levels in the same period^45^, we suspect that absolute TEE levels may have also increased but that relative EE levels may have decreased in line with the observed associations with such indicators in our study.

In conclusion, age, sex and body composition are main biological determinants of human energy expenditure. Results from this nationally representative sample using gold-standard methodology may serve as reference values for other population studies.

## Supporting information

NDNS EE Supplement

## Acknowledgements

We are grateful to all study participants who gave their time to contribute to the NDNS. The survey was jointly funded by Public Health England and the UK Food Standards Agency. There are many people who have been involved in DLW measurement and its interpretation; we wish to thank the following (listed in alphabetical order) at the MRC Dunn Nutrition Unit, and/or at MRC Human Nutrition Research: Darren Cole, Andy Coward, Janet Day, Peter Davies, Kevin Donkers, Rachel Elsom, Abdollah Ghavami, Gail Goldberg, Cheryl Kidney, Alison Lennox, Marilena Leventi, Owen Mugridge, Melanie Nester, Sonja Nicholson, Elise Orford, Ann Prentice, Janine Rattke, Malcolm Sawyer, Priya Singh, Ivonne Solis-Trapala, Toni Steer, Birgit Teucher, Antony Wright. In addition, we thank Beverley Bates and all staff involved in NDNS from NatCen for coordinating and conducting the field work.

The authors were supported by the UK Medical Research Council (unit programme numbers. MC_UU_12015/1, MC_UU_12015/3, U105960371) and the NIHR Biomedical Research Centre in Cambridge (IS-BRC-1215-20014). TL was funded by the Cambridge Trust.

The authors contributed to the present manuscript as follows: Idea for analysis (SB); acquisition, analysis or interpretation of isotope data (MV, LB, PP); epidemiological data analysis (SB, TL); drafting of the manuscript (SB, TL); revising work critically for important intellectual content (all authors); approval of the final version before submission (all authors except LB). There are no conflicts of interest.

