## Supplementary material for "Descriptive epidemiology of energy expenditure in the UK: Findings from the National Diet and Nutrition Survey 2008 – 2015": NDNS EE Supplement

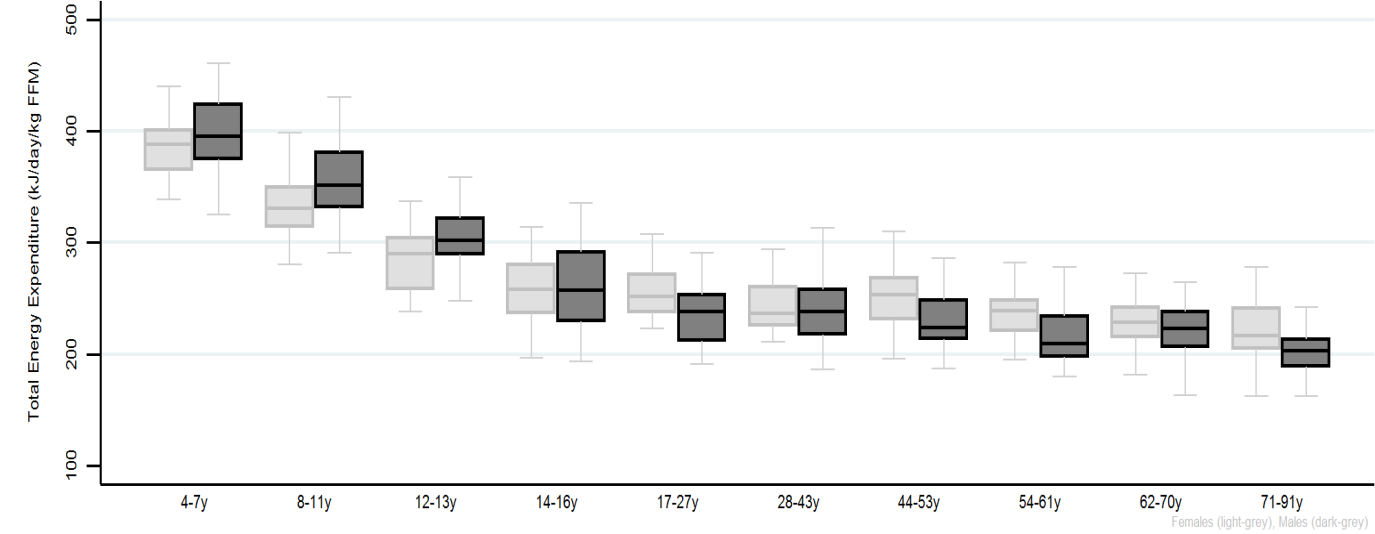

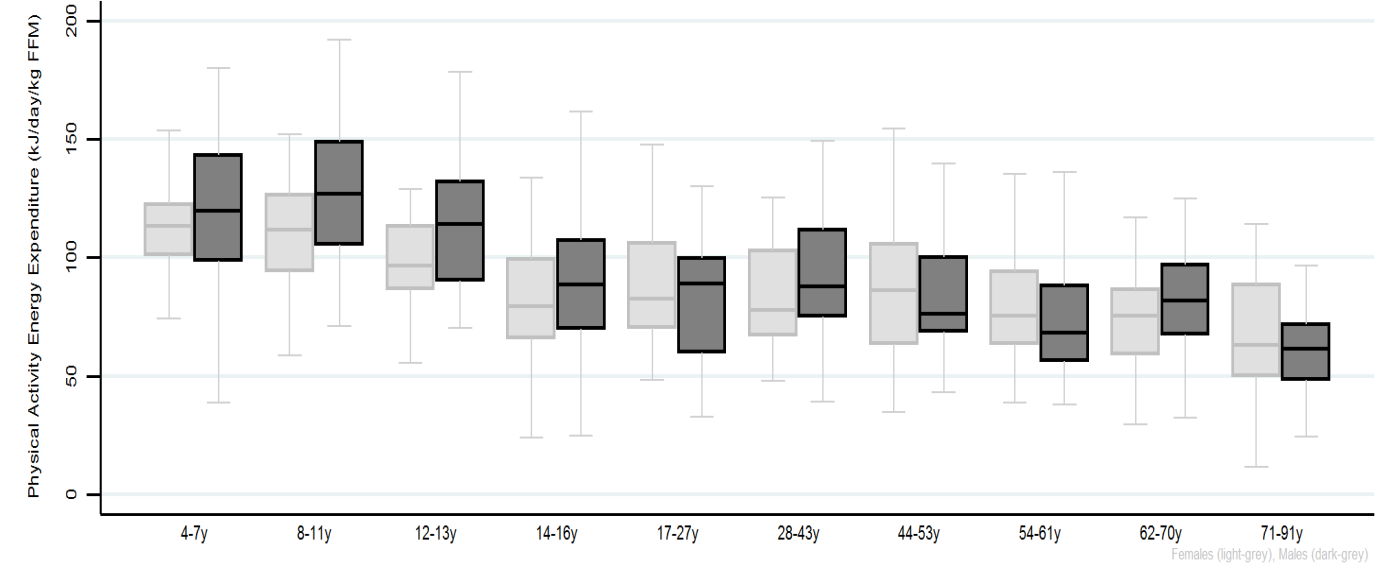

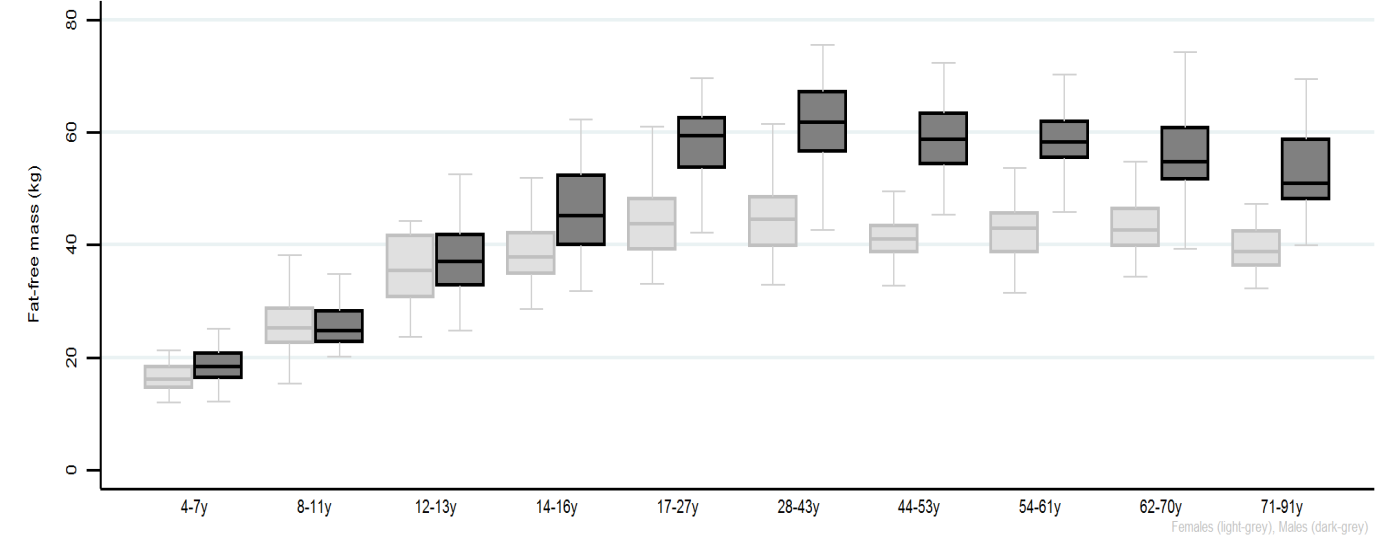

**Figure S1**. Total and Physical Activity-related Energy Expenditure per kg fat-free mass by age (approximate deciles) and sex groups (Females= light grey, Males= dark grey). Bottom panel shows stratified fat-free mass.

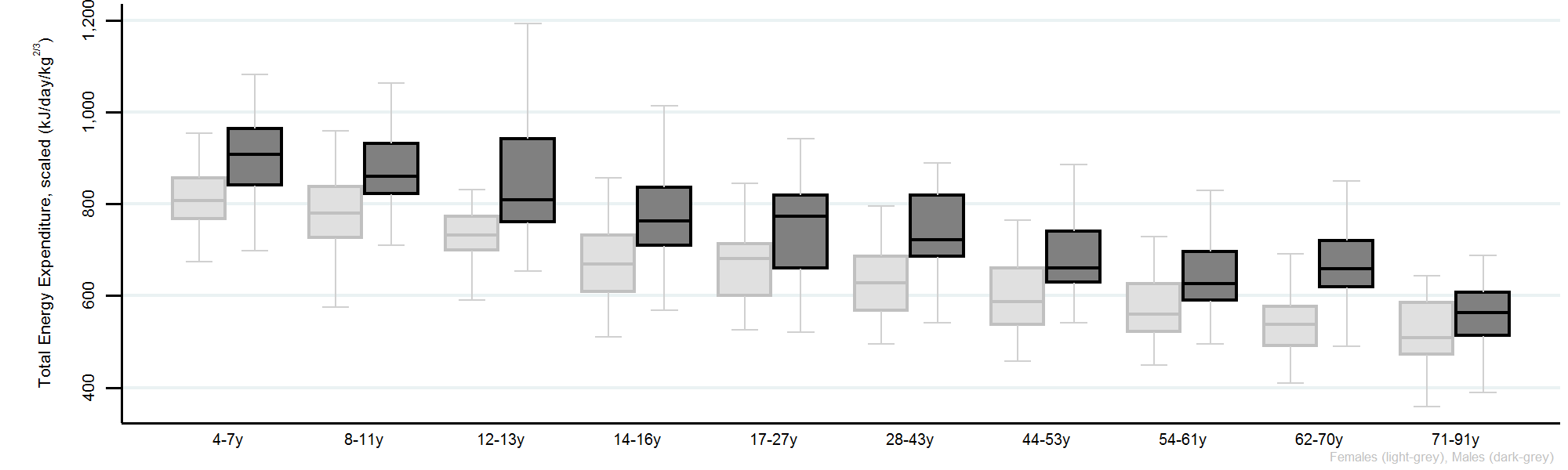

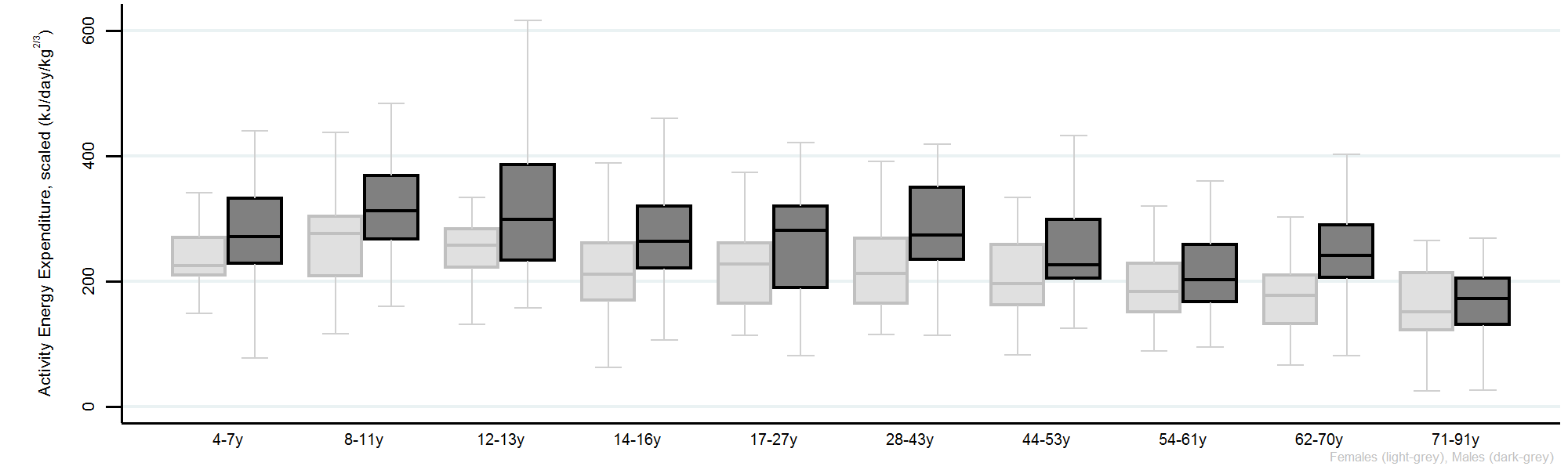
**Figure S2**. Allometrically scaled Total and Physical Activity-related Energy Expenditure per kg^2/3^ total body mass by sex groups (Females= light grey, Males= dark grey) and age (approximate deciles).

**Supplement Table S1:** Sensitivity analysis modelling PAEE per kg fat-free mass from stratifying variables (mutually adjusted)

| **Outcome: PAEE kJ/ day / kg FFM** |  |  | |  |  |
| --- | --- | --- | --- | --- | --- |
|  | **Women** | | | **Men** | |
|  | PAEE  (kJ / day / kg FFM) | | C.I. | PAEE  (kJ / day / kg FFM) | C.I. |
| **Model 1 (FMI Category)** |  | |  |  |  |
| **Age** |  | |  |  |  |
| 4-10y | Reference | |  | Reference |  |
| 11-15y | -16.38*** | | -24.66; -8.11 | -9.52* | -19.06; 0.03 |
| 16-49y | -23.89*** | | -31.54; -16.24 | -36.22*** | -45.18; -27.25 |
| 50-64y | -27.03*** | | -35.24; -18.82 | -43.91*** | -53.22; -34.60 |
| 65-91y | -38.78*** | | -47.21; -30.36 | -55.82*** | -65.87; -45.77 |
| **Year of Study** |  | |  |  |  |
| 2008-2011 | Reference | |  | Reference |  |
| 2012-2015 | 0.59 | | -4.47; 5.64 | -1.49 | -7.57; 4.59 |
| **Season** |  | |  |  |  |
| Spring | 1.27 | | -2.24; 4.78 | 5.14** | 1.08; 9.21 |
| Winter | 1.88 | | -1.73; 5.50 | 0.35 | -4.05; 4.76 |
| **Region** |  | |  |  |  |
| South England | Reference | |  | Reference |  |
| North England | -1.59 | | -7.36; 4.18 | -0.68 | -7.68; 6.32 |
| Scotland, Wales, Northern Ireland | 5.59* | | -0.90; 12.07 | 3.79 | -4.13; 11.71 |
| **FMI Category** |  | |  |  |  |
| 1st Tertile | Reference | |  | Reference |  |
| 2nd Tertile | -4.49 | | -10.69; 1.70 | -4.00 | -11.22; 3.22 |
| 3rd Tertile | -10.70*** | | -17.18; -4.21 | -8.68** | -16.00; -1.36 |
| Constant | 113.32*** | | 106.09; 120.55 | 126.95*** | 117.82; 136.08 |
| **Model 2 (BF% instead of FMI category)** |  | |  |  |  |
| F: <30% M: <25% | Reference | |  | Reference |  |
| F: 30-40% M: 25-35% | -7.06* | | -14.28; 0.16 | -6.13* | -13.43; 1.17 |
| F: >40% M: >35% | -17.08*** | | -24.79; -9.37 | -10.40** | -18.91; -1.90 |
| 95% confidence intervals in parentheses |  | |  |  |  |
| *** p<0.01, ** p<0.05, * p<0.1 |  | |  |  |  |
